# Pairwise graph edit distance characterizes the impact of the construction method on pangenome graphs

**DOI:** 10.1101/2024.12.06.627166

**Authors:** Siegfried Dubois, Matthias Zytnicki, Claire Lemaitre, Thomas Faraut

**Affiliations:** Université de Toulouse, INRAE, ENVT, F-31320 Castanet-Tolosan, France; Univ Rennes, CNRS, Inria, IRISA - UMR 6074, F-35000 Rennes, France; Unité de Mathématiques et Informatique Appliquées, INRAE, Chemin de Borde Rouge, F-31320, France

## Abstract

**Motivation:** Pangenome variation graphs are an increasingly used tool to perform genome analysis, aiming to replace a linear reference in a wide variety of genomic analyses. The construction of a variation graph from a collection of chromosome-size genome sequences is a difficult task that is generally addressed using a number of heuristics. The question that arises is to what extent the construction method influences the resulting graph, and the characterization of variability.

**Results:** We aim to characterize the differences between variation graphs derived from the same set of genomes with a metric which expresses and pinpoint differences. We designed a pairwise variation graph comparison algorithm, which establishes an edit distance between variation graphs, threading the genomes through both graphs. We applied our method to pangenome graphs built from yeast and human chromosome collections, and demonstrate that our method effectively characterizes discordances between pangenome graph construction methods and scales to real datasets.

**Availability:** *pancat compare* is published as free Rust software under the AGPL3.0 open source license. Source code and documentation are available at https://github.com/dubssieg/rs-pancat-compare.

**Supplementary information:** Supplementary data are available online at https://doi.org/10.5281/zenodo.10932490. Code to replicate figures and analysis is available online at https://github.com/dubssieg/pancat_paper.

## 1 Introduction

One of the primary aims of genetics is to analyze how genetic variability impacts phenotype variability. In the standard approaches of genomic analysis, we use the concept of reference sequence: a fully-resolved, high quality assembly which stands for a golden standard for a species.

However, a single sequence cannot encapsulate the whole diversity of a population; nested variations and complex structural variants cannot be positioned over a reference genome and represented as a table with confidence [1]. Rather than considering each of these genomes individually, the aim of the pangenomic approach is to combine the information from multiple genomes. It is the core of the pangenomic approach, which aims to aggregate multiple genome sequence information, improving the quality of read mapping and variant genotyping [2, 3]. To aggregate this information, a data structure, the variation graph, has been introduced. It is a sequence graph where each genome is embedded as a path in the graph with the consecutive nodes corresponding to successive segments on the associated genome sequence [4]. By its construction, variations emerge from the graph topology. In such structures, shared sub-paths correspond to shared genomic regions between genomes and divergent paths to variations, whether small or large, such as inversions, insertions, deletions and substitutions.

The faithful representation of variability is a key to any downstream analysis. Variant genotyping using variation graphs relies heavily on the structural analysis of the graph, including features like bubbles [5]. To ensure accurate reporting of variants, the graph must faithfully represent the variations through the arrangement of nodes. If this is not achieved, variants may be reported inaccurately or not at all. Additionally, a poorly constructed graph can induce errors, creating false variants. A significant portion of the quality is derived from the genomes used, but the choices made by the algorithms also substantially impact the final results. Construction methods for these structures are relatively recent, with *state-of-the-art* pangenome builders such as Minigraph-Cactus [6] and PGGB [7] receiving frequent updates. Inducing a variation graph from a collection of chromosome-size genome sequences poses a significant challenge that is generally addressed through various heuristics, starting with the choice of an alignment algorithm. Moreover, as there is no exact description of the ideal properties for a variation graph [8], various approaches for criteria to optimize have been developed.

In the current literature, variation graphs are described mainly with simple metrics such as the number of nodes and edges. Pangenome graph builders use different alignment algorithms, do not have the same rules for graph induction and apply different post-processing steps. Recent studies have shown that using multiple tools on the same input data produces different graphs [9, 10, 11], however there is no method to quantify or qualify the differences between them. Recently, a method to compare the contents of two graphs using elastic-degenerate strings [12] was published, which addresses the question of the difference of sequence content between graphs, but not how the same sequences are differently embedded in the graph.

Pairwise graph comparison is a well-studied topic, with various measures and algorithms [13] for general purpose graphs that rely vastly on topology. Variation graphs are particular graphs that are made of labelled nodes representing parts of the genomes. Those labels and the way they are scattered in the graph structure convey meaningful information about the potential conservation or variation of sequences, which must be considered when comparing graphs.

In the work presented here, we aim to characterize the differences between variation graphs derived from the same set of genomes. To pursue this objective, we present an algorithm to compute a pairwise segmentation distance between variation graphs, exploiting the way the genomes are embedded in the graph structure. Our method not only provides a metric to evaluate the extent to which two graphs differ but also the ability to pinpoint where differences are, enabling to analyse their distributions in the graph as well as to locate them at the nodes. Then, we apply this method to real pangenome graphs to compare graphs built from two state-of-the-art pipelines, and assess their differences. Lastly, we investigate these differences — their impact, position, and size — in order to provide insights into their impact on pangenome analysis.

## 2 Materials and methods

### 2.1 Sequence graphs and genome segmentation

Let Γ = {Γ_0_, Γ_1_, … Γ_*n*_} be a set of genomes. A genome Γ_*i*_ is represented by a string *w*_1_ … *w*_*m*_ with *m* being the size of the genome, on the alphabet {*A, C, G, T*}.

This genome collection Γ can be represented by a sequence graph *G*(Γ) = (𝒱, ℰ). This structure is a directed graph where each node *v* ∈ 𝒱 is labelled by a word *w*, that exists in at least one genome of Γ, in forward or reverse orientation. Any edge *e* ∈ ℰ links two vertices whose labels are contiguous in at least one genome. Edges are directed and annotated at both ends, conveying both reading direction and order of connected nodes.

A pangenome graph 𝒢 (Γ) = (𝒱, ℰ, 𝒫) is a sequence graph extended by a collection of paths 𝒫. A path consists of an oriented and ordered list of vertices linked by edges in the graph. A path represents a continuous input sequence, be it a scaffold or a chromosome; meaning a single genome can be embedded in the graph by numerous paths; for the sake of simplicity, we will consider that each genome is represented as a single path although our method can be generalized to encompass can be generalized to graphs with genomes split into several chromosomes or scaffolds. We require the pangenome graphs to be *complete*. We say that a pangenome graph is *complete* if the sequence of each element of Γ can be read by following one path from the path collection 𝒫 (Definition 1).

#### Definition 1 (Complete pangenome graph)

*A complete pangenome graph* 𝒢 (Γ) = (𝒱, ℰ, 𝒫) *is a pangenome graph where* |𝒫| = |Γ|, *and every genome of* Γ *is represented in exactly one path of* 𝒫. *Concatenation of every label associated with the vertices of* 𝒫_*i*_, *in respect with their orientation, is exactly the sequence of* Γ_*i*_.

By the definition of a pangenome graph, the successive nodes encountered in the traversal of path 𝒫_*i*_ correspond to consecutive genomic intervals along the genome Γ_*i*_ associated to this path. We can consider a path as an ordered and oriented list of contiguous genomic intervals on a genome. This describes a *segmentation* of each genome in the graph, where genomic intervals defined by the traversed vertices are separated by *breakpoints*.

#### Definition 2 (Breakpoint)

*A breakpoint b is a position in a genome where the graph structure breaks the continuity between two consecutive genomic intervals. The existence of a breakpoint on a genome* Γ_*i*_ *is associated with an edge between the two vertices supporting the two associated labels in the path* 𝒫_*i*_. *This position is expressed in number of basepairs from the start of the genome* Γ_*i*_.

We define ℬ_*i*_ = *b*_0_ … *b*_*n*_ as the set of breakpoints on Γ_*i*_ induced by 𝒢. This set of breakpoints correspond to a segmentation of Γ_*i*_, which reflects the evolutionary relationship with the other genomes in Γ but also heavily depends on the graph construction process. In order to compare the graphs, we can compare how differently the same genomes are segmented in different graphs, resulting in comparing pairwisely breakpoint sets for each genome.

We propose in the following paragraphs an algorithm which compares two pangenome graphs on the behalf of the pairwise comparison of breakpoint sets associated to each genome of Γ. Let 𝒢g_*a*_(Γ) = (𝒱^a^ ℰ^a^, 𝒫^a^) and 𝒢_*b*_(Γ) = (𝒱^b^ ℰ^b^, 𝒫^b^) be two complete pangenome graphs sharing the same set of genomes Γ. The idea of our comparison method is to find differences between the two segmentations of Γ_*i*_ by uncovering the smallest set of breakpoints that differentiates 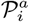 and 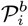.

We define two reciprocal operations, *merge* and *split*, which correspond respectively to the suppression and addition of a breakpoint in a path at a position *x*. Thus, *merge*(*x*, 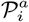) removes the breakpoint at the position *x* in the path 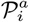, while *split*(*x*, 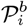) adds on the path 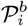 a breakpoint at the position *x*. Merges and splits are particular breakpoints that are missing in 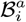 or in 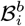. If we consider 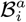 and 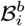 as two sets, the symmetric difference of the breakpoints of the two segmentations is the union of all required merges and splits, and we call the size of this set the segmentation distance (Definition 3). Summation of all segmentation distances between two graphs gives the distance between them (Definition 4).

#### Definition 3 (Path segmentation distance)

*The segmentation distance d*_*s*_ *for a genome* Γ_*i*_ *present in two graphs* 𝒢_*a*_ *and* 𝒢_*b*_ *is the minimum set of operations enabling to transform one segmentation into the other. It corresponds to the breakpoints that are exclusive to one of the breakpoints sets* 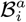 *and* 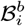.

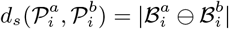

*Where x* ⊖ *y is the symmetric difference between x and y*.

#### Definition 4 (Graph segmentation distance)

*The segmentation distance d between two graphs containing the same genome set* Γ *is the summation of all the segmentation distances over the genomes of* Γ:

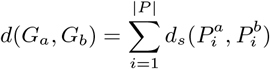

We can generalize this definition to any pair of graphs that shares at least one common genome, by computing the intersection of paths of the two graphs and applying the segmentation distance computation only on the intersection.

### 2.2 Algorithm

For each genome Γ_*i*_, we browse linearly and simultaneously through the two associated segmentations to find the specific breakpoints between the two breakpoint lists. In Algorithm 1, 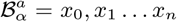 and 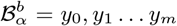 are breakpoints of two paths 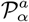 and 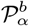 representing Γ_*α*_, such that both paths represent the same sequence but can differ in segmentation. We consider at each step a position *p* in the string Γ_*i*_, which increases at each iteration to the next closest breakpoint across both segmentations. ℳ is the set storing merges, and 𝒮 the set storing splits.

#### Algorithm 1 Distance *d*_*s*_ over a single path

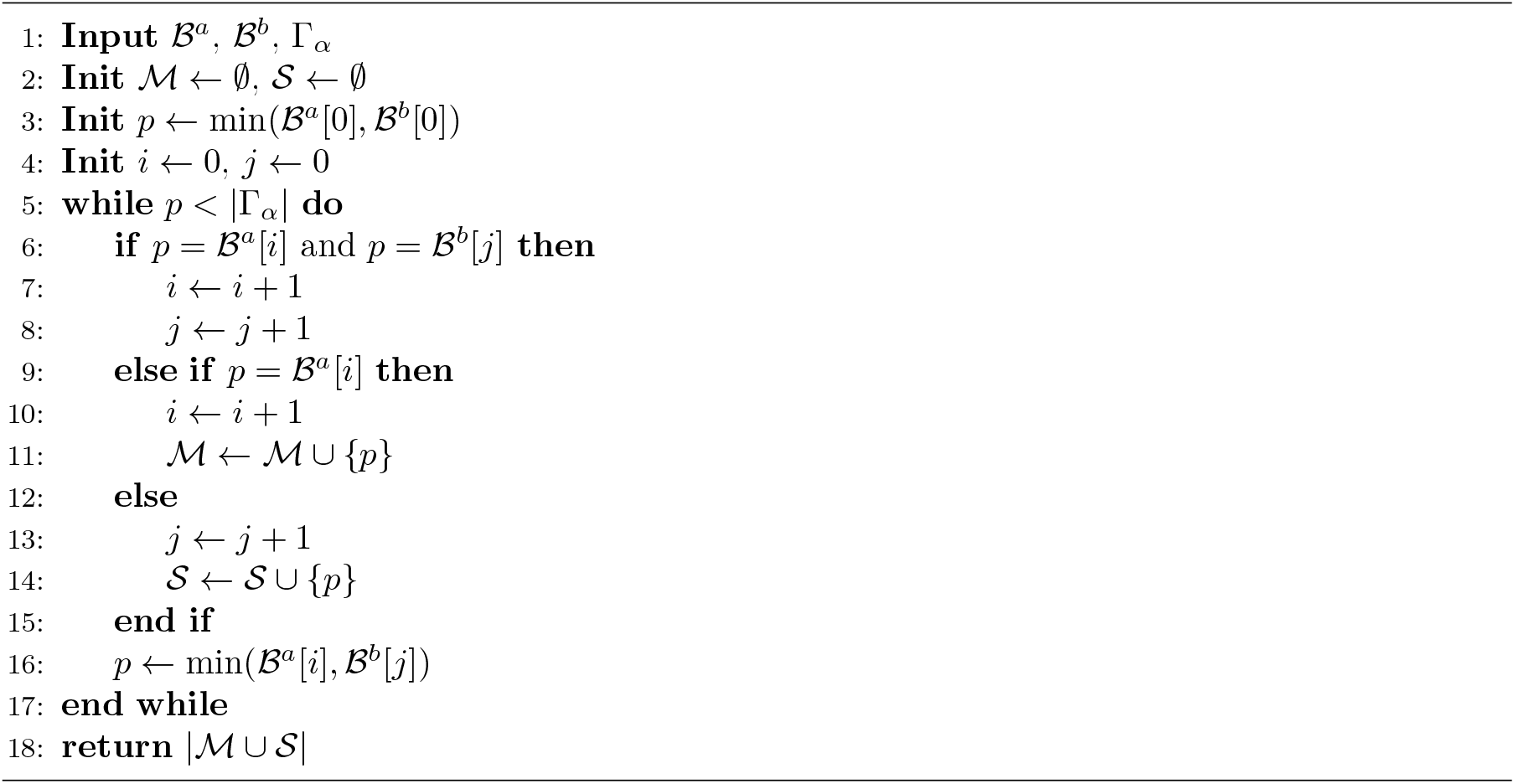

With our definition of editions, each operation applies on a single breakpoint, by adding it or removing it in the other segmentation. The number of differing breakpoints across both segmentations is the summation of the breakpoints that are specific to any of the two segmentations: ℬ^*a*^ ⊖ ℬ^*b*^. When we iterate, we take into account every breakpoint (ℬ^*a*^ ∪ ℬ^*b*^) but we only count as edition breakpoints that are not common to both segmentations (ℬ^a^ ∪ ℬ^b^ \ ℬ^a^ ∩ ℬ^b^), which is the symmetric difference between the two sets, thus ensuring that our algorithm gets the minimal number of editions. Algorithm 1 executes for a genome Γ_*i*_ with two breakpoints sets 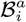 and 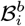 in a total of 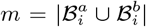 steps. It results in a complexity of 𝒪(*m*) for the path segmentation distance. When a segmentation distance between two graphs is computed, as we execute this algorithm once per genome, we end up with a complexity of 𝒪 (*n* · *m*) with *n* being the number of genomes in Γ and *m* the size of the union of the breakpoints of the two segmentation of each genome.

We now give the intuition that our measure is a segmentation distance. The following properties are given at the path level. As the sum of distances are a distance, if *d*_*s*_ is a segmentation distance then *d* is too:

- *Positive*: our metric is a summation of the number of elements of two sets, ℳ and 𝒮. Those counts can be positive or zero (meaning segmentation are identical), and from Algorithm (1), having zero merges and zero splits means that we have the same segmentation on both paths, thus validating 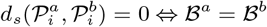
- *Symmetry* : merge and split are reciprocal operations. From Equation 3, if we find the minimal couple of sets ⟨ℳ^*a*^, 𝒮^*a*^⟩, we can define the reciprocal couple ⟨ℳ^*b*^, 𝒮^*b*^⟩ where ℳ^*b*^ = 𝒮^*a*^ and 𝒮^*b*^ = ℳ^*a*^, thus granting symmetry as we keep the same number of total operations.
- *Triangle inequality* : let 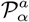, 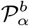 and 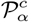 be three paths in three distinct graphs representing a same genome Γ_*α*_. We need to show that 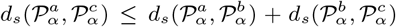. It exists for each segmentation a set of breakpoints ℬ. We can transform our equation into 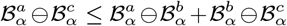, which is true from the definition of the symmetric difference.

### 2.3 Case study

The algorithm was applied both on published pangenome graphs, the HPRC human draft pangenome graphs [10], and graphs build from telomere-to-telomere yeast assemblies [14].

For the yeast dataset, we selected chromosome 1 from 15 samples and constructed the graphs using the Minigraph-Cactus pipeline [6] (*mgc*, v2.9.0) and the PanGenome Graph Builder [15] (*pggb*, v0.6.0). From the selected yeast assemblies, we built multiple graphs, varying the reference sample for *mgc*and the order of secondary genomes. We also constructed a corresponding *pggb* graph with the same assemblies, ensuring that all comparisons were made between graphs representing the same input data exactly. All graphs were verified as complete pangenome graphs.

Graphs built with *mgc* had the *clip* and *filter* parameters set to zero, ensuring that the entire sequence was embedded in the structure. Both tools were run with all other parameters kept at their default settings. The graphs did not undergo any post-processing, and graphs from *pggb* were collected before the *smoothxg* step to ensure completeness. We identified genomic variants in these graphs using *vg deconstruct* [5] (v1.56.0).

In this study, for the purpose of comparing graphs, we make a distinction between “shared variants” and “private variants”. A “shared variant” is defined as a variant that has the same position, reference allele, and alternate alleles in both graphs (ignoring the order of alternates). In contrast, a “private variant” refers to any variant that does not meet these criteria. Tandem repeats on the linear reference genome are computed using *TandemRepeat Finder* [16] (v4.09.1) and are expressed as the number of bases per kilobase. Unique *k*-mers are computed using a simple sliding window and are expressed as the number of different *k*-mer per kilobase.

For the human dataset, we analyzed differences between *mgc* and *pggb* graphs from the HPRC dataset (year 1) for chromosomes 1 and 21 individually. We computed editions between the full *mgc CHM13* graph (built with mgc v2.6.4, HPRC year 1, v1.1), without clipping or filtering, and the *pggb* graph (built with pggb v0.2.0+531f85f, HPRC year 1, v1.0). Variants were extracted from the raw CHM13 VCFs supplied along the pangenome graphs. Human pangenome graphs from the HPRC can be challenging to compare due to differences in scaffold sets. Not all genomes included in these graphs are telomere-to-telomere, thus implying that chromosomes may be composed of multiple scaffolds. The assignment of these scaffolds to chromosomes was performed using distinct methods for the two tools, resulting in graphs that include both shared sequences and tool-specific sequences (see Supplementary Table 1). In this case, the distance is computed on the intersection of scaffolds of the two graphs.

**Table 1:** Datasets used to evaluate the comparison tool. Minigraph-Cactus = *mgc*, PanGenome Graph Builder = *pggb*. Org: Organism. Chr: chromosome number. #hap: haplotype count in graph. Length: length of the reference genome. Total length: sum of all node lengths in graph. Variants: Variants are called against the haplotype that was used as reference to build the *mgc* graph. Note that the number of variants is the only metric that is not graph-level in this table. Merges and splits are operation counts computed in the direction from the *mgc* graph to the *pggb* one.

| Org. | Chr. | #hap. | Length (bp) | Tool | Total length (bp) | Nodes | Edges | Variants | Merges | Splits |
| --- | --- | --- | --- | --- | --- | --- | --- | --- | --- | --- |
| yeast | 1 | 15 | 222,424 | *mgc* | 386,083 | 34,889 | 48,130 | 9,213 | 35,956 | 44,097 |
|  |  |  |  | *pggb* | 350,140 | 27,213 | 37,755 | 8,626 |  |  |
| human | 21 | 90 | 45,090,682 | *mgc* | 423,514,455 | 2,541,744 | 3,486,748 | 574,574 | 12,949,063 | 79,480,033 |
|  |  |  |  | *pggb* | 273,835,317 | 2,760,531 | 3,882,969 | 841,419 |  |  |
| human | 1 | 90 | 248,387,328 | *mgc* | 1,449,752,909 | 6,951,299 | 9,620,356 | 2,224,237 | 32,362,430 | 189,702,647 |
|  |  |  |  | *pggb* | 1,117,392,094 | 11,109,656 | 15,398,037 | 3,456,614 |  |  |

Yeast graphs and supplementary data are available at https://doi.org/10.5281/zenodo.10932490.

### 2.4 Implementation

All editions between graphs are computed using *rs-pancat-compare* (v0.1.2), which is our Rust implementation of Alg. 1 available at https://github.com/dubssieg/rs-pancat-compare. After storing a mapping of the nodes to their respective length, we do a parallel reading of the node lists described in the path and asynchronously iterate on both paths to compute each operation. The output is a tab-separated file containing one edition per line described by its path name, its position in the path, a one-letter encoding of the operation, nodes on which the operation applies to in first and second graph, and individual ending breakpoints for both nodes. Results are subsequently processed using Jupyter Notebooks, which are available at https://github.com/dubssieg/pancat_paper.

## 3 Results

### 3.1 Comparing graphs

We applied our method to two datasets: yeast graphs built from 15 telomere-to-telomere assemblies of chromosome 1 [14] and graphs of chromosome 1 and 21 from the HPRC human draft pangenome [10]. We computed metrics for both the Minigraph-Cactus (*mgc*) and the PanGenome Graph Builder (*pggb*) versions of the graphs and measured the distances between them (Table 1).

#### 3.1.1 Differences between pangenome graphs

For the yeast dataset, the graphs of the chromosome 1, constructed from the same 15 genome set with *mgc* and *pggb* show significant differences for every metric (Tab. 1). The *pggb* graph is smaller in all aspects, containing 27,213 nodes (compared to 34,889 for *mgc*) and a total length of 350 kb (compared to 386 kb). The length of a graph is computed by summing the length of all the labels of its nodes. Representing the same genome content with a reduced length can result from a failure of one of the graph to summarize shared genome content or a tendency of the other graph to adopt a more compact representation, for example for the repsentation of repeated sequences, resulting in an increased number of cycles in the genome paths. This is a known difference between *mgc* and *pggb*, with the latter constructing more compact graphs at the expense of a larger number of cycles [9]. These differences are also reflected in the variant sets reported by *vg deconstruct*, with 9,213 variants identified in the *mgc* graph and 8,224 in the *pggb* graph. Among these variants, 6,291 are shared between the two graphs, 2,922 are unique to the *mgc* graph, and 1,933 are unique to the *pggb* graph. For the same chromosome, analysis of the operations needed to transform the *mgc* graph into the *pggb* graph reveals that 44,907 splits and 35,956 merges are needed. By dividing by the total number of characters in the genome collection, these numbers represents an average of 25.24 editions per kilobase.

For the human dataset, the observed metrics follow the same trends. The density of editions also remains within a similar range: the chromosome 21 graph shows an average of 25.33 editions per kilobase, while chromosome 1 displays approximately 10.22 editions per kilobase. It has to be noted that the two graphs do not share exactly the same set of scaffolds. For each HPRC graph (*mgc* or *pggb*), the scaffolds included in the pangenome graph for each chromosome is defined by the construction strategy used that apparently differed (see Supplementary Table 1). This might impact the segmentation of the other scaffolds in the graphs and possibly of the segmentation of the shared scaffolds which will be captured by our algorithm and identified as graph discrepancies. These scaffolds, however, represents a small fraction of the whole scaffold set (6.24% for the chromosome 21 and 1.84% for the chromosome 1). We believe therefore that this does not change the global picture and that the observed differences result essentially from the construction algorithms rather than from the difference in the initial set of scaffolds.

By analyzing the positions of the editions, we can calculate metrics such as their lengths, providing insight into the importance of the differences. Editions can be projected as splits onto one graph or the other (see Methods). The locations of edition breakpoints within the nodes reveal how the graphs differ in their representation of genome segments. Here, the size of an edition is defined as the length of the smallest segment resulting from the split on the node where the edition occurs, regardless of other editions affecting the same node. The median edition size is notably large (yeast chromosome 1: 8 bp; human chromosome 21: 114,247 bp; human chromosome 1: 21,477 bp), which can be attributed to the prevalence of large nodes (nodes longer than 50 bp represent 67.12%, 95.69%, and 95.99% of the base pairs of the respective graphs). Some large nodes can exhibit a very large number of editions. A single node, for example, of the yeast *mgc* chromosome 1 graph has to be edited 2,339 times to match the corresponding nodes in the *pggb* graph. In a similar manner, the human *mgc* chromosome 21 graph has a node that has 1,354,801 editions, and the human *mgc* chromosome 1 graph has a node that has 3,660,206 editions. It seems that those are nodes that were left non-aligned by one tool but where the other one forced the alignment. When distinguishing editions based on the length of the nodes they affect, small nodes (≤ 50 bp) consistently exhibit a median, upper, and lower quartile edition size of 3 bp across all graph pairs analyzed. Small nodes do not account for the majority of editions (32.82% for yeast chromosome 1, 8.32% for human chromosome 21, and 12.74% for human chromosome 1), yet editions affecting these nodes are still over-represented relative to their coverage in the respective graphs. There is a bias towards the tip of the nodes regarding edition density, and also on the number of editions that affects the smallest nodes (see Supplementary Figure 1).

**Figure 1:**
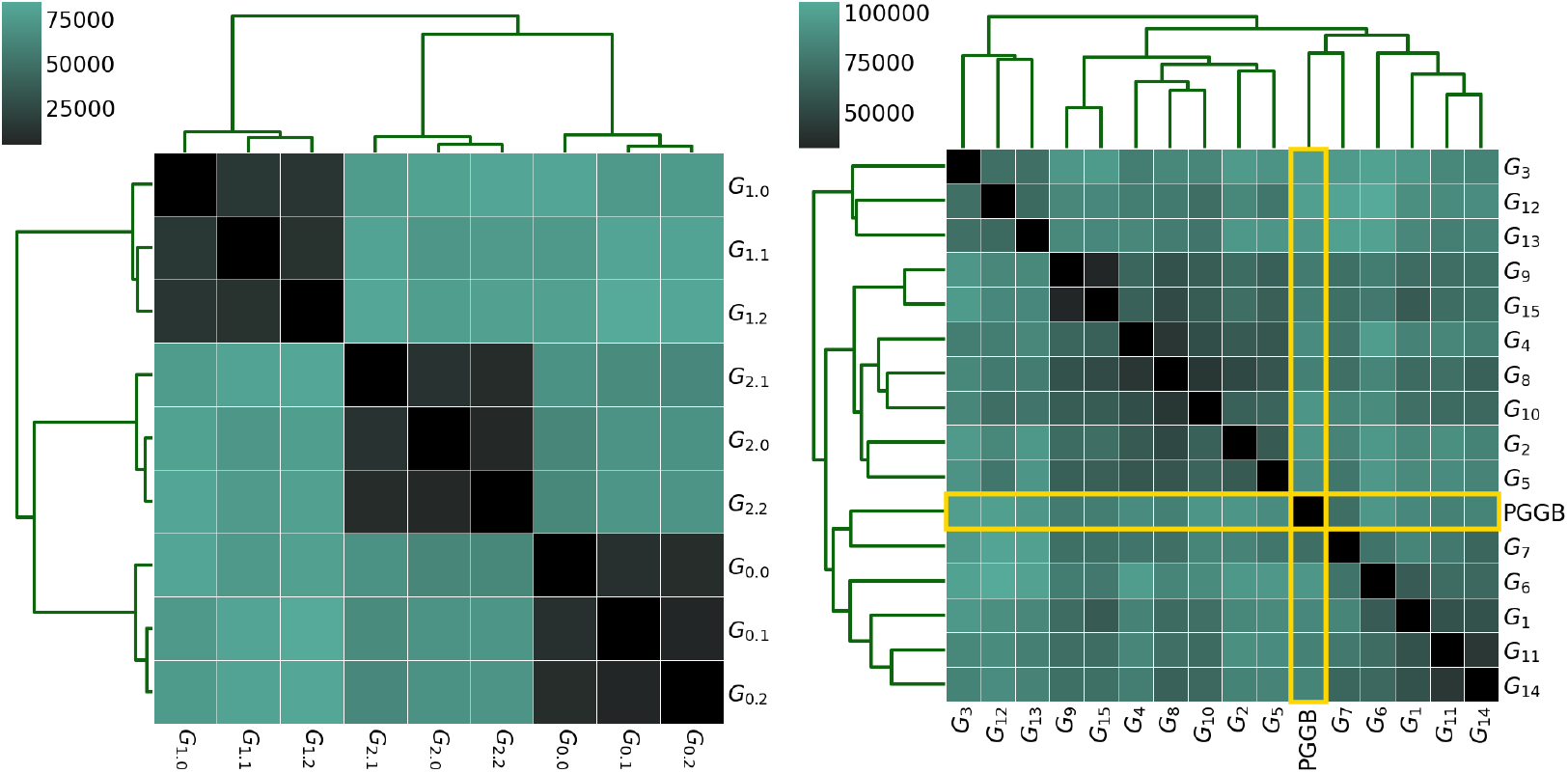
Hierarchical agglomerative clustering of distances between graphs, computed with *pancat*. A lower distance means that less breakpoints needs to be added or removed to form the other graph. A) Comparison of 9 *mgc* graphs; each triad uses a shared reference and only differs by secondary genomes order. B) Comparison of 15 *mgc* graph and 1 *pggb* graph; *mgc* graphs differs by the genome used as reference.

#### 3.1.2 Impact of the reference choice

The *mgc* pipeline adds incrementally genomes atop of a chosen reference. We applied our method to compute segmentation distances between graphs built with different reference genomes and input genome orders. According to the edition metric, in *mgc*, changing the order of secondary genomes has less impact than switching the reference genome (Figure 1.A). Graphs that share the same reference genome cluster very well, with editions counts between 5k and 15k, whereas distances between graphs with different reference genomes range from 65k to 80k editions.

When comparing graphs made with *mgc* featuring different references to the *pggb* graph that does not take any reference nor genome order, it appears that changing the reference in *mgc* can have more impact on segmentation distance than switching tools. Graphs generated with *mgc* do not cluster separately from the *pggb* graph with the same genomes. The mean distance between the *pggb* graph and *mgc* graphs is about 88k, while for *mgc* graphs it ranges from 70k to 89k on average. For almost every *mgc* graph we built, there exists another *mgc* graph that has a higher distance to the first than to the *pggb* graph. This confirms the distance clustering (Figure 1.B), as the *pggb* graph is not an outlier in terms of distance from the multiple *mgc* graphs.

This observation has significant implications for graph analysis, as the differences directly affect the number of shared variants and in turn the number of private variants observed between graphs. Private variants — those that differ in their detection or representation across graphs — are closely correlated with the number of editions (Spearman r=0.82, p-value=4.43e-64) whereas it is not the case for shared variants (Spearman r=-0.07, p-value=3.00e-01). This strong correlation with private variants supports the idea that segmentation distance serve as a robust metric for identifying areas featuring differently reported variants (see Supplementary Figure 2).

**Figure 2:**
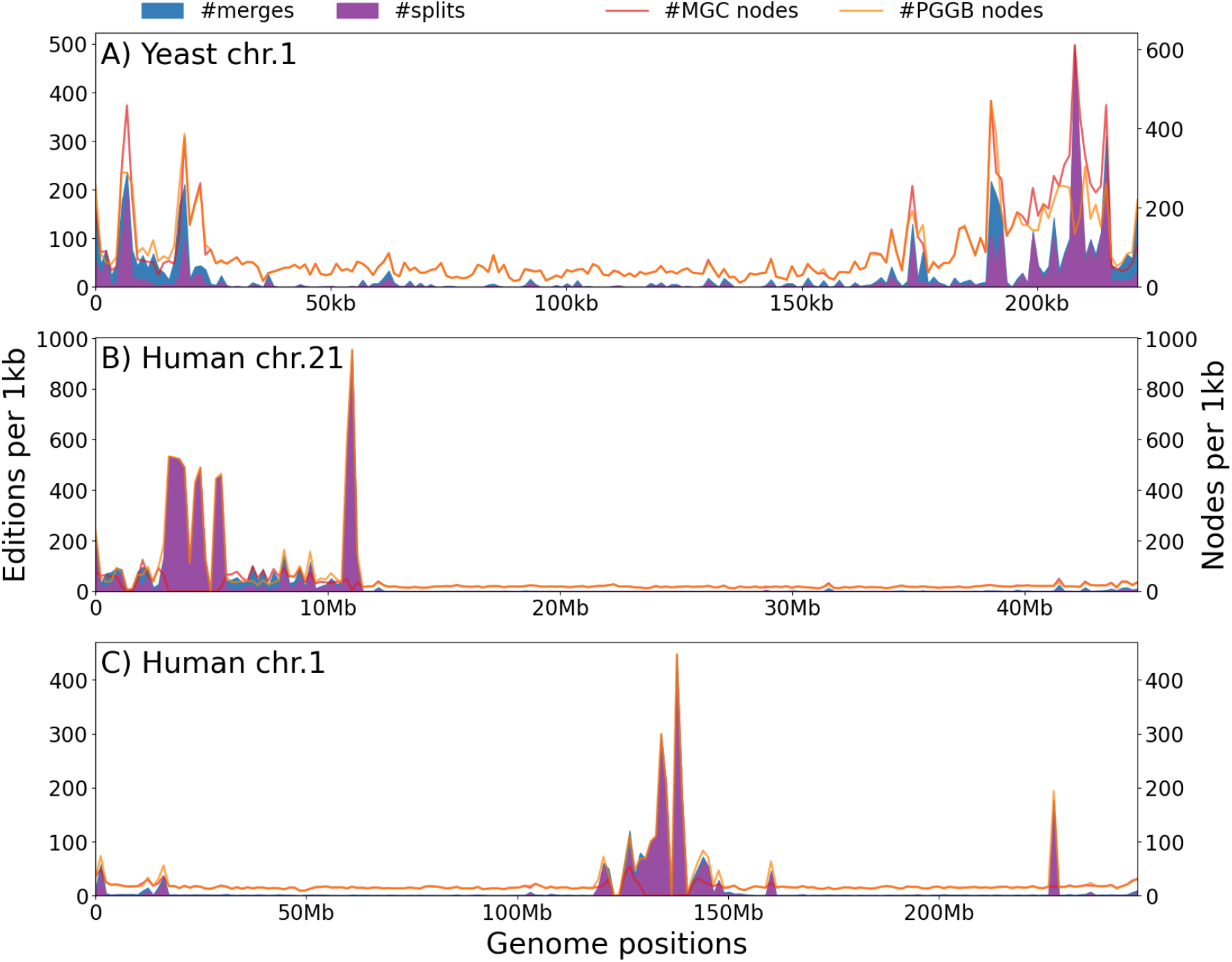
Projections of editions on a linear genome. In such projections, only editions affecting this particular path in the graph are displayed, as well as for the node counts. Projections highlight the existence of edition hotspots. A) Yeast, chromosome 1, projected on CASBJU01. B) Human, chromosome 21, projected on CHM13. C) Human, chromosome 1, projected on CHM13.

### 3.2 Distribution along the genome

The segmentation distance can be measured between graphs, offering a metric to compare the way genomes are embedded in the graph. Going further, the precise locations of the editions along a given genome provide clues about the causes of graph discrepancies. However, these editions can be distributed evenly along the genomes, acting like a noise. We will show that it is not the case, and link these editions to genome properties.

#### 3.2.1 Edition hotspots

Projected along a genome, editions are not evenly distributed, but rather concentrated in some regions, creating hotspots of differences (Figure 2). This behavior is consistent across all datasets, with peaks on centromeric or sub-telomeric regions (see Supplementary Table 2). The same pattern is observed when changing the reference for graph construction, as well as when projecting onto other haplotypes within the same graph.

**Table 2:** Comparison of timings and peak memory usage over diverse datasets. Peak memory takes into account the overhead of loading graphs in memory. Results are collected from jobs executed on a single core of a 13th Gen Intel® Core™ i7-1365U @ 3.6GHz. Timings and memory consumption were recorded using *heaptrack*.

| Organism | chromosome | wall time | RAM<sub>peak</sub> |
| --- | --- | --- | --- |
| yeast | 1 | 0m01s | 3.0 MB |
| human | 21 | 5m08s | 383 MB |
|  | 1 | 17min42s | 1.2 GB |

#### 3.2.2 Relations between genome composition and graph

The structure of the graph reflects the relationship between genomes but, as previously mentioned, is also influenced by the specific choices of the construction process, in particular the way genomes are aligned. This structure has an impact on downstream analysis, and we can expect that the editions, which reflect the differences between graphs, can locally be in line with differing number of nodes or differently reported variants.

Edition and node numbers in the same window are correlated for yeast (Spearman r=0.71, p-value=3.58e-63 for chromosome 1) as well as for human (r=0.63, p-value=2.33e-45 for chromosome 21 and r=0.51, p-value=5.13e-28 for chromosome 1). As splits create nodes and are the most common operation type, this result is not surprising. Investigating variants reported for *mgc* and *pggb* for yeast allows to separate variants into shared or private. In chromosome 1 of yeast, 56.44% of reported variants are common, 26.22% are private to *mgc* and 17.34% to *pggb* (Tab. 1). There is a correlation between the number of variants and the number of editions that is more pronounced for private variants, as expected (Spearman’s r=0.97 for private variants, against r=0.41 for shared ones on yeast chromosome 1, and respectively for human chromosome 21 and 1 r=0.93 and 0.91 for private variants, r=0.19 and -0.02 for shared variants, (Supplementary Figure 3). Hence, the correlation between editions and private variants is not only at graph-scale but also at the local scale of a genome we are calling the variants against.

**Figure 3:**
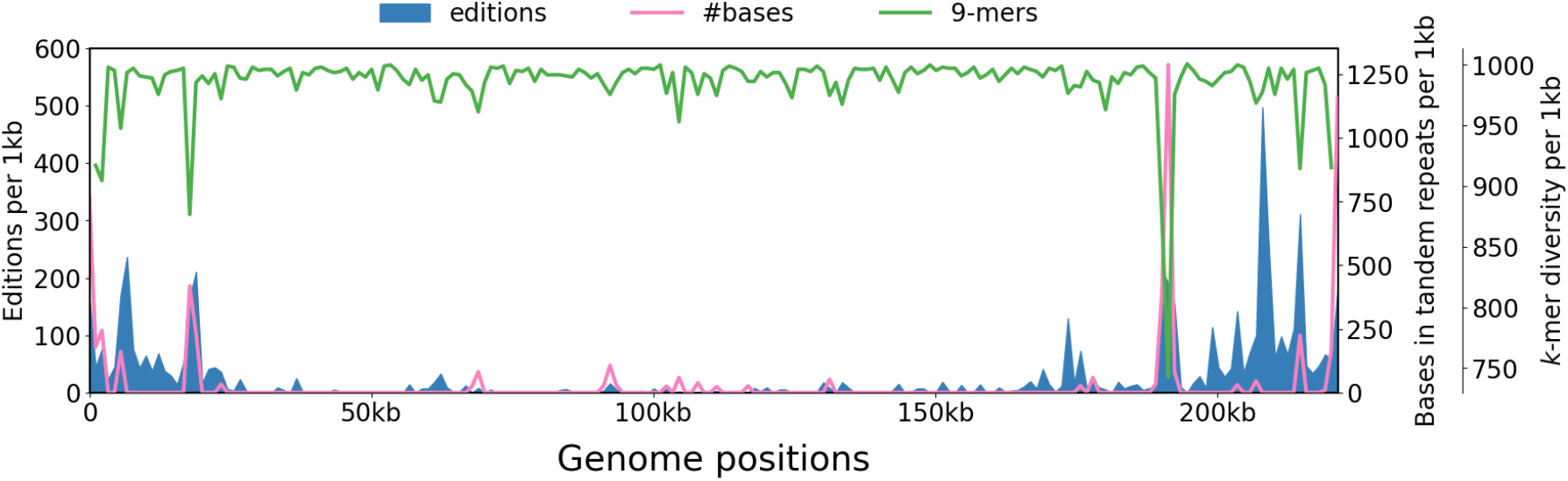
Projections of distinct *k*-mers and tandem repeats on a linear yeast genome. Tandem repeats are expressed as the number of bases involved in tandem repeat over a 1kb window, and distinct *k*-mers as the number of unique 9-mers in a 1kb window. Editions are expressed as the summation of splits and merges.

Investigating the relationship between genome composition and edition distribution reveals strong local correlations between genome complexity and edition hotspots. Distinct *k*-mers (number of unique 9-mers in a sliding window) and tandem repeats (number of bases reported as tandem repeats in a sliding window) are strongly correlated to the number of editions for some peaks. This is particularly true for the peaks around 20kb and 190kb for the yeast chromosome 1 graph (Figure 3), but genome features does not explain everything (peaks around 220kb for yeast chromosome 1 could not be explained with either of these features). Nevertheless, these features remain valuable indicators. For yeast, the correlation is moderate (Spearman r=0.35, p-value=2.71e-07 for tandem repeats and r=0.40 with p-value 5.06e-09 for unique 9-mers) but is stronger for human (respectively for chromosome 21 and 1, Spearman r=0.79 and 0.79 with p-value=2.71e-43 and 8.63e-44 for tandem repeats, r=0.46 and -0.25 with p-value 3.73e-11 and 3.09e-4 for unique 9-mers; see Supplementary Figure 4). This highlights the influence of genomic features on graph construction, particularly in regions that are challenging to align, where the process becomes significantly more complex.

### 3.3 Scalability

Our implementation scales reasonably well, with the comparison of graphs of human chromosomes with 90 haplotypes from the HPRC ranging from 5min to 18min (respectively for chromosome 21 and 1) using at most 1.2 GB for the human chromosome 1 (Table 2). With such numbers, we estimate one can compare the entire human pangenome built with *mgc* and *pggb* with 24 cores and 30 Gb of RAM in under 30 minutes. The running time is related to the length of the paths, in nodes, and RAM growth is conditioned by the number of nodes in both graphs.

## 4 Discussion

In this work, we presented a novel method to compute a distance between a pair of variation graphs, that allows for the establishment of a segmentation-related dissimilarity metric and pinpoint where those differences are located. We defined an edition distance between pangenome graphs as the sum of segmentation differences across common genomes that are embedded in both graphs.

We think that understanding how graphs differ is a key step to improve graph construction tools as well as for qualification of accurate variants. Variants are reported by current state-of-the-art graph-based variant detection tools through the enumeration of topological patterns called bubbles [5].

Starting and ending positions of variants correspond to breakpoints, and the genome segmentation which result from the alignment determine the graph structure. In this work, we showed that regions that are difficult to align lead to differences in graph topology, which in turn affect the detection of variants within the graph. The density of editions along a genome can be interpreted as a confidence indicator for variants called against this genome, though our tool cannot determine which representation is optimal. This highlights a significant challenge in pangenomics: defining objective functions for graph construction and establishing methods to effectively represent complex variation sites within the graph.

This work emphasizes the importance of the construction method choice. In Minigraph-Cactus, the choice of the reference genome is crucial, as it forms the backbone of the entire graph. Opting for PGGB instead of Minigraph-Cactus also influences the resulting graph. Swapping the reference genome in Minigraph-Cactus or using PGGB results in a similar order of magnitude for the number of editions between the graphs, which also translates into differences in private variants. As a community, we lack a clear understanding of the best practices for pangenome graph construction, and there is a dire need for methods to assess both the quality of the graph and its ability to accurately summarize or alter the information derived from the individual genomes.

This distance definition does not capture the full picture of the differences between two graphs. Two graphs may share an identical segmentation but have different topologies. For example, a label shared by multiple haplotypes could be duplicated in one graph, creating bubbles that could be simplified. Similarly, graphs might express the same segmentation but either form a cycle or duplicate a series of labels. Depending on what we want to measure, we may need to define new operations, such as node duplication and fusion, to account for these differences. This would result in an hybrid distance that balances topology and segmentation.

One of the questions that arises is how graph normalization would impact this distance. Some edition peaks are related to tandem repeats and low-complexity regions. While those factors does not explain all the differences, a significant part of our editions are confined to the tips of the largest nodes or to small nodes, which can be thought as alignment choices that might be resolved through a normalization process. In the context of the representation of the majority of indels, left-normalization is widely used, and may be a satisfactory way to mitigate this issue. To our knowledge, no such tool or algorithm currently exists for graphs, but standardizing variation graphs could be a good way to eliminate biases and ensure consistent results from any graph built from the same data [17].

Our distance metric provides insights into the genome breakdown within the graph, and we extended our analysis to propose hypotheses regarding these differences. Although we explored features that could explain graph differences, we do not have precise answer on good practices to build a pangenome graph. Building pangenome graphs remains a complex task, that requires a careful choice of the genomes that will make the backbone of the graph. It also requires a critical assessment of the pangenome graph builders, to ensure the building process is made with respect to the desired representation of variants. With different existing building methods, we hope that our work can further help identify low-confidence variants and facilitate the investigation of variation representation in pangenome graphs, and open discussions on pangenome graph benchmarking and quality assessment.

## Competing interests

No competing interest is declared.

## Author contributions statement

S.D., C.L., T.F. and M.Z. conceived the experiments, S.D. conducted the experiments, S.D., C.L., T.F. and M.Z. analysed the results. S.D., C.L., T.F. and M.Z. wrote and reviewed the manuscript.

## Acknowledgments

We acknowledge Benjamin Linard for his insights during the preliminary work, as well as Sandra Romain for the discussions about the representation of variation inside pangenome graphs. We thanks the GenOuest bioinformatics core facility for providing the computing infrastructure. This work was supported by state funding managed by the French National Research Agency under the France 2030 program [grant number ANR-22-PEAE-0005]. A CC-BY public copyright license (https://creativecommons.org/licenses/by/4.0/) has been applied by the authors to the present document, in accordance with the grant’s open access conditions.

## Notes

https://github.com/dubssieg/rs-pancat-compare

https://github.com/dubssieg/pancat\_paper

https://doi.org/10.5281/zenodo.10932490

### Competing Interest Statement

The authors have declared no competing interest.

