## Supplemental Tables and Figures for "Pairwise graph edit distance characterizes the impact of the construction method on pangenome graphs"

### Supplementary Materials

Siegfried Dubois[1,2], Matthias Zytnicki[3], Claire Lemaitre[2], and Thomas Faraut[1]

[1] *GenPhySE, Université de Toulouse, INRAE, ENVT, 31320 Castanet-Tolosan, France*
[2] *Univ Rennes, CNRS, Inria, IRISA - UMR 6074, F-35000 Rennes, France*
[3] *Unité de Mathématiques et Informatique Appliquées, INRAE, Chemin de Borde Rouge, F-31320, France*

### Supplementary Table 1

**Table S1:** Graphs from the HPRC year 1 created from *minigraph-cactus* and the *PanGenome Graph Builder* did not use the same scaffold attribution method, resulting in graphs with a majority of sequences shared in-between them, but also with some private parts. It is important to underline that the choices weren't made by the pangenome builders but during the data curation. As our method focuses on segmentation distances, we could not compare haplotypes that were present in one graph and not in another. Is it hard to tell to which extent the presence and absence of those scaffolds in the graphs ended up changing the topology of the graph, however we can still use the yeast graph as a ground standard as it was build to offer a fair comparison between the two graph builders, using strictly the same data.

| | Yeast, chr1 | Human, chr21 | Human, chr1 |
|---|---|---|---|
| Total *mgc* size | 3,172,121 | 3,770,658,781 | 21,735,427,417 |
| Total *pggb* size | 3,172,121 | 4,012,006,987 | 22,552,620,745 |
| Shared size | 3,172,121 | 3,648,504,353 | 21,734,915,199 |
| Private *mgc* size | 0 | 122,154,428 | 512,218 |
| Private *pggb* size | 0 | 363,502,634 | 817,705,546 |

### Supplementary Figure 1

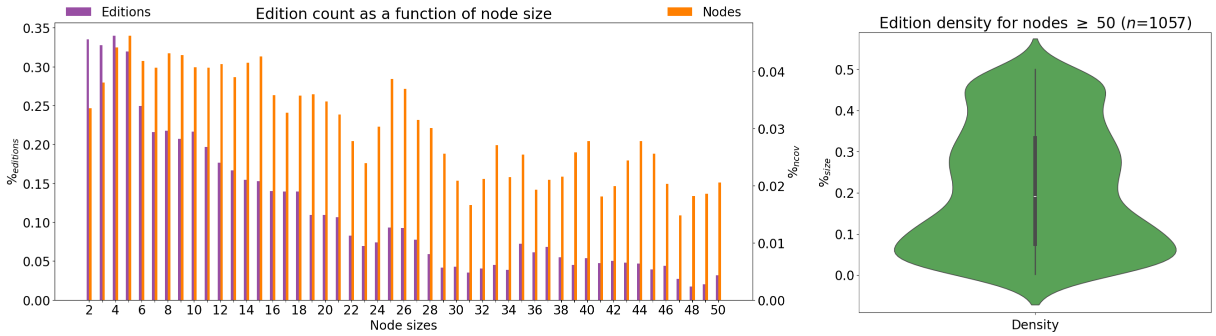


**Figure S1:** Distribution of edition sizes over the nodes of the graph. A) For every node size $n$ between 2 and 50, number of editions (normalized by node length) and number of nodes of size $n$. The longer a node is, the lower the number of edited positions is. B) Edition position as a percentage of size for long nodes ($\geq$50 bp). We can observe that more editions are located near the tips of the node than in the middle.

### Supplementary Figure 2

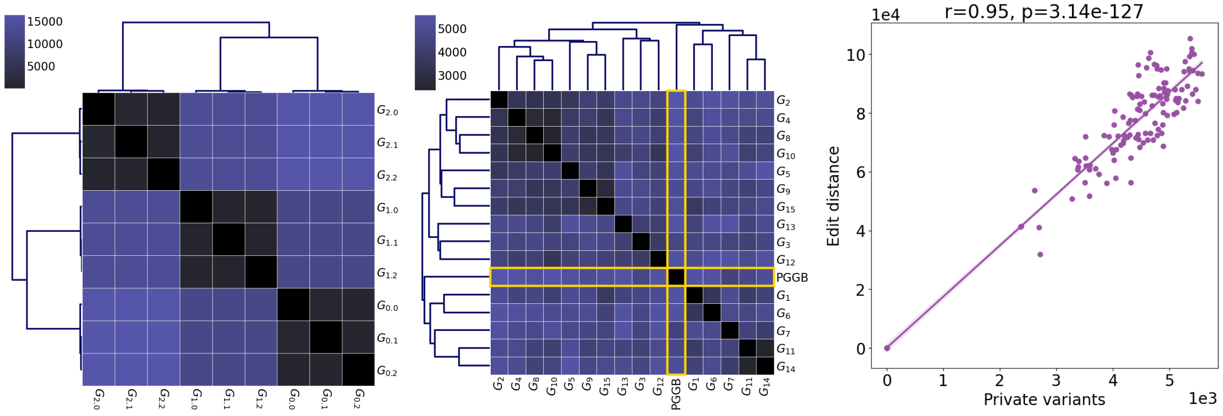


**Figure S2:** Hierarchical agglomerative clustering of private variants between yeast graphs. The compared graphs are the same as the one from Fig. 1 of the main manuscript. A lower count means that less variants are not in common between two graphs. All variants have been called against the same reference. Clustering of private variants ends being really similar to the clustering of editions.

### Supplementary Table 2

**Table S2:** Highest edition peaks annotations across yeast and human data. Annotations are accessed *via* the UCSC genome browser. $\epsilon$: Editions. ORF: Open Reading Frame. TR: Tandem Repeat. cenSat: Centromeric Satellite. periSat: Peri-Centromeric Satellite.

| Org. | Chr. | $\epsilon/kb$ | Median (in $\epsilon/kb$) | Peak position (in Mb) | Peak (in $\epsilon/kb$) | UCSC annotation |
|---|---|---|---|---|---|---|
| yeast | 1 | 25.24 | 5.39 | 0.208 - 0.209 | 497 | dubious ORF + multiple TRs |
| | | | | 11.0 - 11.3 | 955 | cenSat |
| human | 21 | 25.33 | 2.00 | 10.8 - 11.0 | 575 | cenSat |
| | | | | 3.2 - 3.4 | 533 | periSat/cenSat |
| human | 1 | 10.22 | 1.63 | 138.0 - 139.0 | 448 | cenSat |

### Supplementary Figure 3

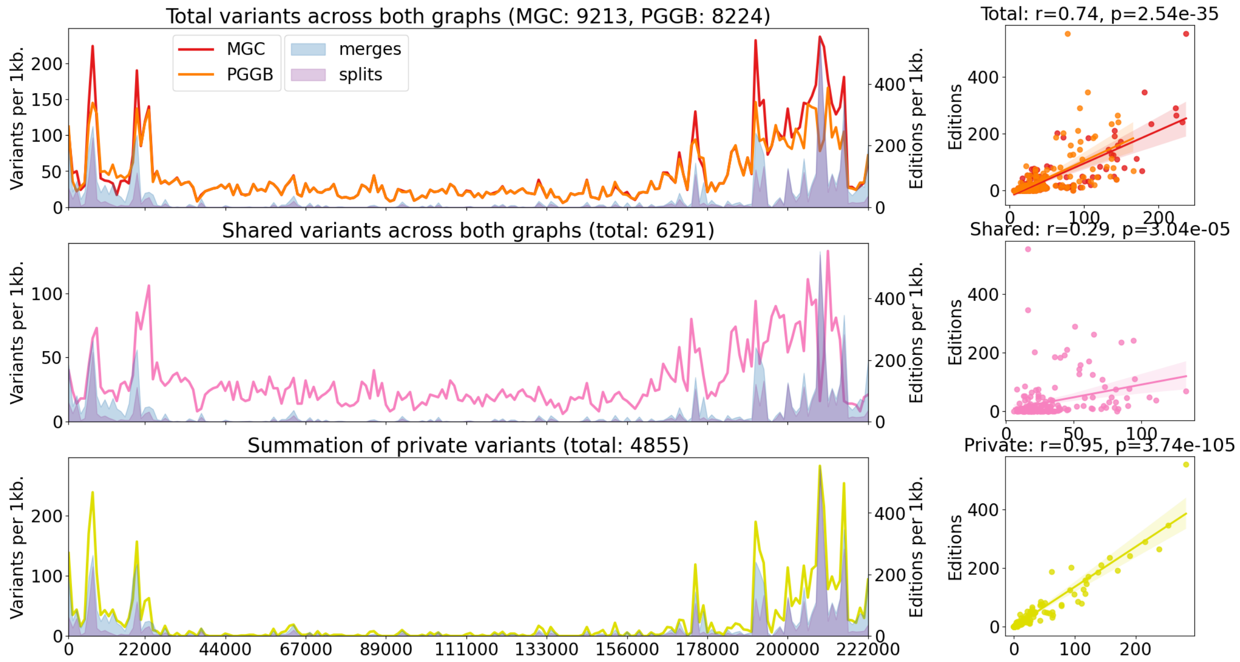


**Figure S3:** Projection along the reference genome of the editions between yeast graphs from *Minigraph-Cactus* and *PGGB*. Variants are computed against this same sequence, and we project their position along the genome.

### Supplementary Figure 4

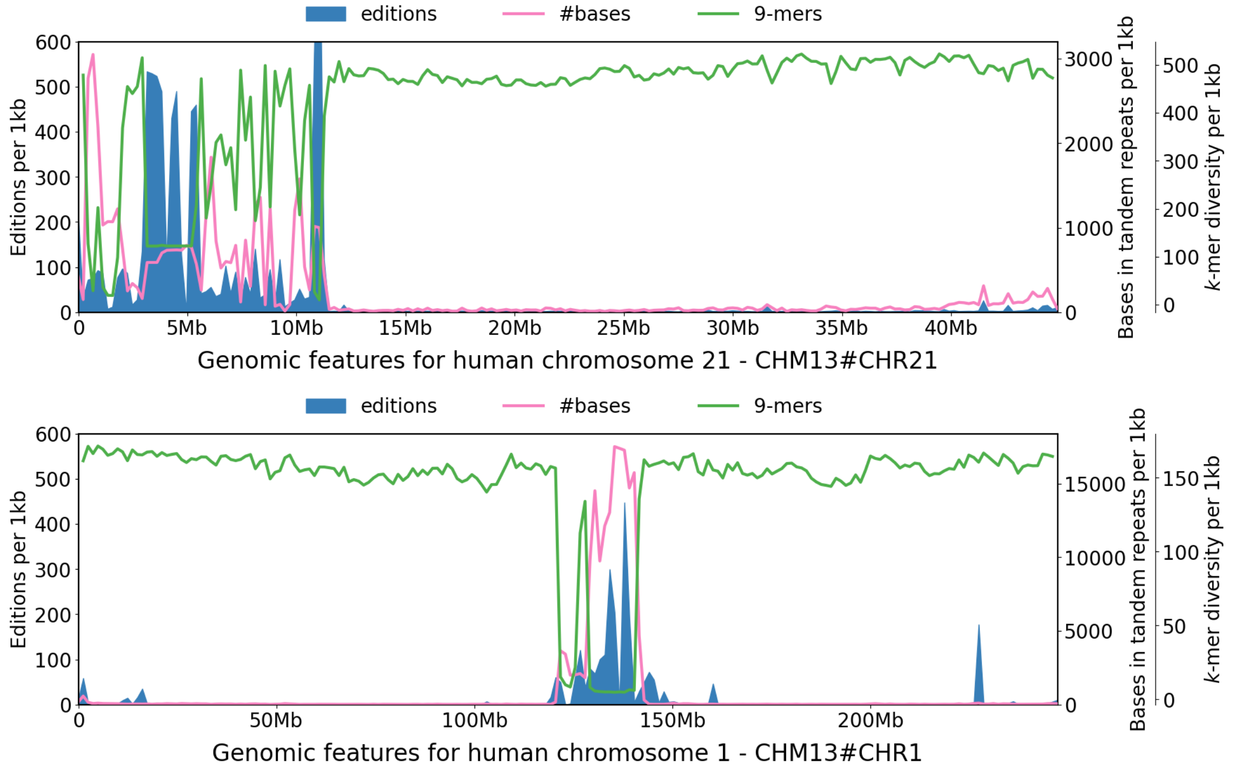


**Figure S4:** Projection along the reference genome of the editions between human graphs from *Minigraph-Cactus* and *PGGB*. Tandem repeats and unique 9-mers are computed on the same genome.
